# Cryo-EM structures of anti Z-DNA antibodies in complex with antigen reveal distinct recognition modes of a left-handed geometry

**DOI:** 10.64898/2025.12.12.693871

**Authors:** Danielle Chin, Yongbo Luo, Yiteng Lau, Nivedita Dutta, Zengyting He, Chaoran Yin, Riley M. Williams, Siddharth Balachandran, Quentin Vicens, Peter Dröge, Dahai Luo

**Author notes:** DHRC and YL contributed equally to this work.

## Abstract

Double-stranded nucleic acids can undergo transitions from canonical B/A-forms to alternate left-handed Z-DNA/Z-RNA (Z-NAs). Z-NAs are implicated in processes such as neuroinflammation in Alzheimer’s disease, Lupus Erythematosus, microbial biofilms, and type I interferon-mediated human pathologies. Since endogenous Z-NA sensors like the Zα domain can induce B-to-Z transitions, monoclonal antibodies (mAbs) Z-D11 and Z22 have been regarded as conformation-specific tools to confirm Z-NA *in situ*, although high-resolution structural information is missing. Here, we employed single-particle cryo-electron microscopy to solve structures of Z-D11 and Z22 bound to synthetic d(CG)_6_ 12mer Z-DNA duplex. Both mAbs form filamentous trimers around the Z-DNA axis, further stabilized by Fab-Fab interactions. The mAbs achieve specificity through extensive contacts to both Z-form backbone strands and the exposed guanine/cytosine bases in the major groove. This mode of recognition is dictated by shape complementarity rather than sequence specificity, sensing the alternating syn/anti backbone torsions and the phosphate zig-zag geometry unique to Z-DNA. Our data also suggest that these mAbs are not inducing B-to-Z transitions under normal physiological conditions. Finally, comparison to other double-stranded NA-binding mAbs defines a similar structural logic adapted to different helical geometry recognition patterns, thus providing a framework for engineering highly specific nucleic acid probes.

## Introduction

Double-stranded nucleic acids (ds-NAs) exhibit remarkable polymorphisms which depend on nucleotide sequence contexts and external conditions. Among prominent examples are transitions from canonical B-/A-form right-handed to left-handed duplex geometries. The resulting alternate conformers, dubbed Z-DNA/Z-RNA (collectively Z-NAs), have been solved at high resolution ^[1, 2]^, and their biophysical and biochemical properties were extensively investigated revealing both substantial similarities as well as important differences ^[3]^.

Following Z-NA discoveries in the 1970s ^[4]^, their biological roles remained elusive for decades ^[5]^. High resolution structures of the Za domain from the dsRNA editing enzyme ADAR1 in complex with Z-NAs eventually demarcated turning points ^[6, 7]^. Experimental evidence now suggests involvement of Z-DNA in microbial biofilms ^[8]^, thymic T cell tolerization ^[9]^, cell death during limb development ^[10]^, extrachromosomal DNA biogenesis ^[11]^, neuroinflammation in Alzheimer’s disease ^[12]^, Lupus Erythematosus ^[13, 14]^, autoimmune photosensitivity ^[15]^, and memory effects in the mouse brain ^[16]^. Z-RNA is being implicated in innate immune reactions following viral infections ^[17-19]^ and Aicardi-Goutière’s Syndrome ^[20]^, cancer immunotherapy ^[21]^, aberrant mRNA splicing ^[22-24]^ and type I IFN-mediated human pathologies ^[25, 26]^.

Za domains and their variants are frequently used in these studies as indicators for intracellular Z-NAs. However, in addition to binding already formed Z-NAs, Za is able to induce B-to-Z transitions in appropriate sequence tracks under physiological conditions, with faster transition kinetics for Z-DNA than Z-RNA ^[27-29]^. Furthermore, Za binds A-form-like duplexes with low micromolar affinity ^[30]^. Hence, monoclonal antibodies (mAbs) raised against Z-DNA have been employed to confirm *in situ* presence of Z-NAs.

The most commonly used anti-Z-DNA mAb is derived from hybridoma clone Z22 ^[31]^. Another mAb employed in early Z-NA studies, Z-D11, exhibited interesting properties such as Z-DNA complex stability half-life of several days, hysteresis, and blockage of *in vitro* transcriptional elongation through Z-DNA templates ^[32-39]^. However, despite their importance in studying the appearance and biological relevance of Z-NAs, detailed structural information for mAb-antigen complexes is still missing.

In the current report, we employed Z-D11 and Z22 mAbs in single-particle cryo-electron microscopy (cryo-EM) and solved their high-resolution Fab structures bound to ds d(CG)_6_ in high salt conditions. We show that both mAbs adopt oligomeric binding modes using extensive contacts to both Z-form duplex backbone strands and the guanine and cytosine bases in the major groove, spanning eight dG:dC base pairs. We also describe the potential influence of d^5^mCdG modification and alternate dGdT:dAdC sequence in Z-conformation on mAb binding via molecular modeling; the latter supported by plasmid-based EMSAs. In addition, we propose that direct Fab-Fab interactions contribute to binding avidity on longer Z-DNA stretches. Finally, *in silico* comparisons with models of mAb S9.6 bound to a right-handed RNA:DNA hybrid and of mAb J2 in complex with ds A-form RNA suggest a shared logic in antibody-NA recognition. Our findings thus point at generalizable principles of ds-NA recognition by antibodies and may help to establish a framework for engineered conformation-specific NA probes.

## Results

### Oligomeric Assembly of Z-D11 and Z22 on Z-DNA

To elucidate Z-DNA recognition modes, we determined the structures of recombinant Z-D11 and Z22 bound to a synthetic 12-mer duplex d(CG)_6_. Complex assembly was optimized using high salt buffer conditions (2.5M NaCl, and 0.7M MgCl2), which induces the B-to-Z structural transition ^[4]^. In fact, original hybridoma screening for Z-D11 was conducted at 4M NaCl using unmodified ^32^P-labeled ds d(CG) oligonucleotides ^[35]^. Z22 hybridoma cells, however, were identified with brominated d(GC) oligonucleotides at physiological conditions ^[31]^.

Analytical size exclusion chromatography (SEC) under these conditions confirmed the formation of large oligomeric complexes for both Z-D11 and Z22, as evident by significant shifts in elution profiles toward higher molecular weights compared to free components (Fig. 1B, 1C). As expected for a structure-specific mAb, Z-D11 showed a substantial free antibody peak under low salt conditions (150mM NaCl) (Fig. 1A). Cryo-EM micrographs revealed distinct assembly pattern: the Z-D11-d(CG)_6_ complex formed long filamentous structures, visualized in both micrographs and 2D classifications (Fig. 1D, 1F), while the Z22-d(CG)_6_ complex appeared as slightly discrete, multi-unit particles (Fig. 1E, 1G).

**Figure 1.**
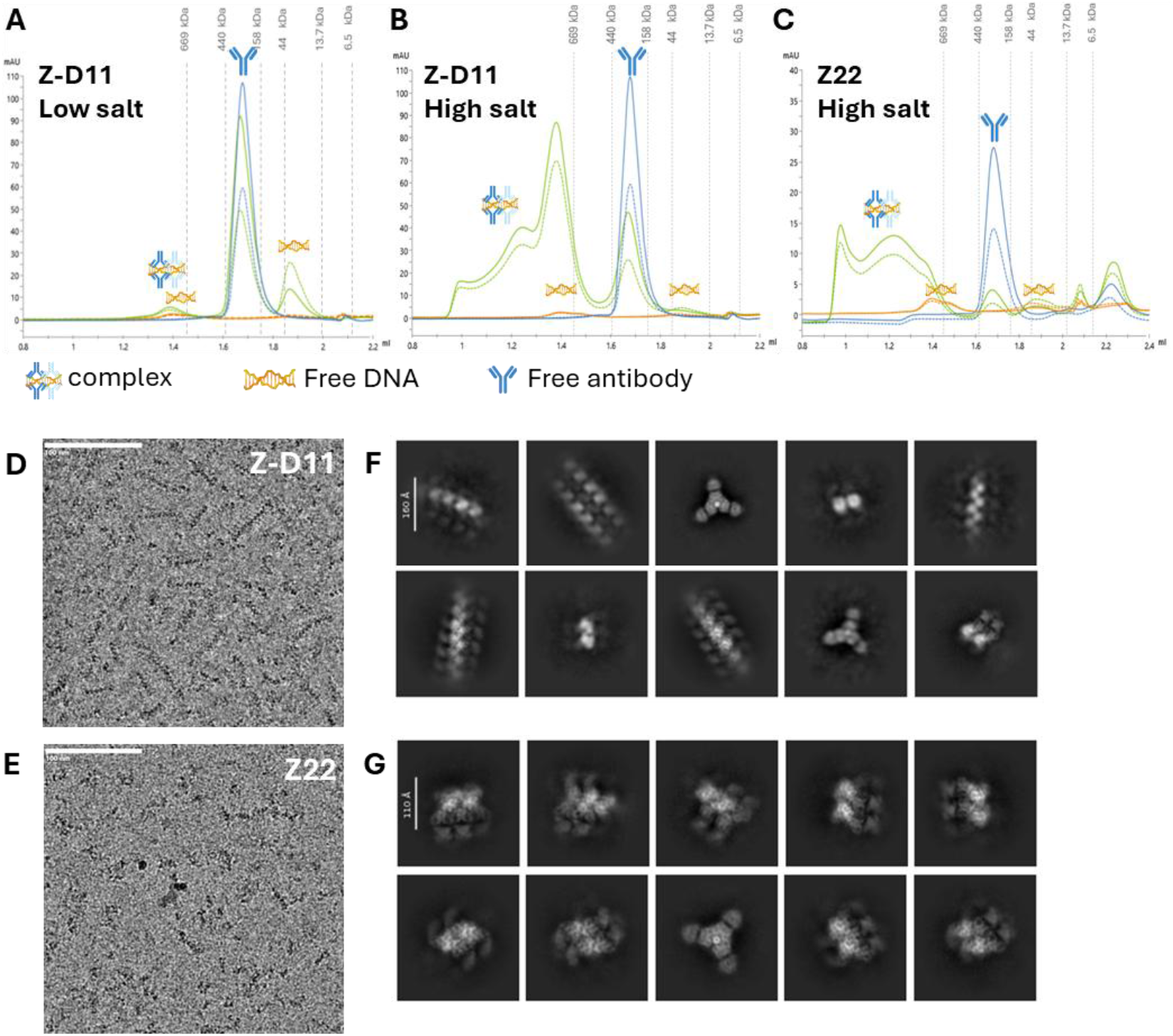
Z-D11 and Z22 mAbs bind to Z-DNA to form filaments. Size exclusion chromatograms of of Z22 in complex with d(CG)_6_ DNA under **A**. low salt and **B**. high salt condition. **C**. Size exclusion chromatograms of Z22 in complex with d(CG)_6_ DNA under high salt conditions. Low salt buffer contains 20mM HEPES pH 7.4, 150mM NaCl; high salt buffer contains 20mM HEPES pH 7.4, 2.5M NaCl and 0.7M MgCl_2_. Representative image of the **D. – E**. cryo-EM micrographs for the respective Z-D11-d(CG)_6_ and Z22-d(CG)_6_ complexes, collected under 165,000x magnification and viewed with 10Å low pass filter. Scale bar represents 100nm. **F. – G**. Representative 2D classes for the respective Z-D11-d(CG)_6_ and Z22-d(CG)_6_ complexes are shown.

### High-Resolution Cryo-EM Structures and Stoichiometry

The final high-resolution structures obtained through single-particle analysis revealed a dimer of trimers assembly (Fig. S1, S2). The Z-D11 complex exhibits an axial alignment along the duplex, with its filamentous nature attributed to extensive contacts between neighbouring Fab heavy chains (Fig. 2A, 2C). Specifically, the Fab-Fab interface in Z-D11 involves heavy chain residues Thr 28 and Gly 26, displaying bond lengths of 3.9 Å (Fig. 2E). For the Z22 complex, initial processing yielded a 2.80Å map showing two trimer units. Utilizing 59,879 particles, further localized refinement upon supplying a mask over the first trimer unit resulted in a 2.67 Å map of a single trimer formation surrounding the Z-DNA (Fig. 2B, 2D). This refined model contains 681 protein residues and 24 nucleotides. The Z22 Fab-Fab interface similarly relies on contacts between heavy chains, involving Asn 28 and Gly 26 with bond lengths ranging from 2.9 Å to 3.7 Å, with additional water-mediated contact between the peptide bonds of the heavy chains. (Fig. 2F). In both structures, the antibodies demonstrate binding that spans eight dG:dC base pairs along the left-handed Z form duplex.

**Figure 2.**
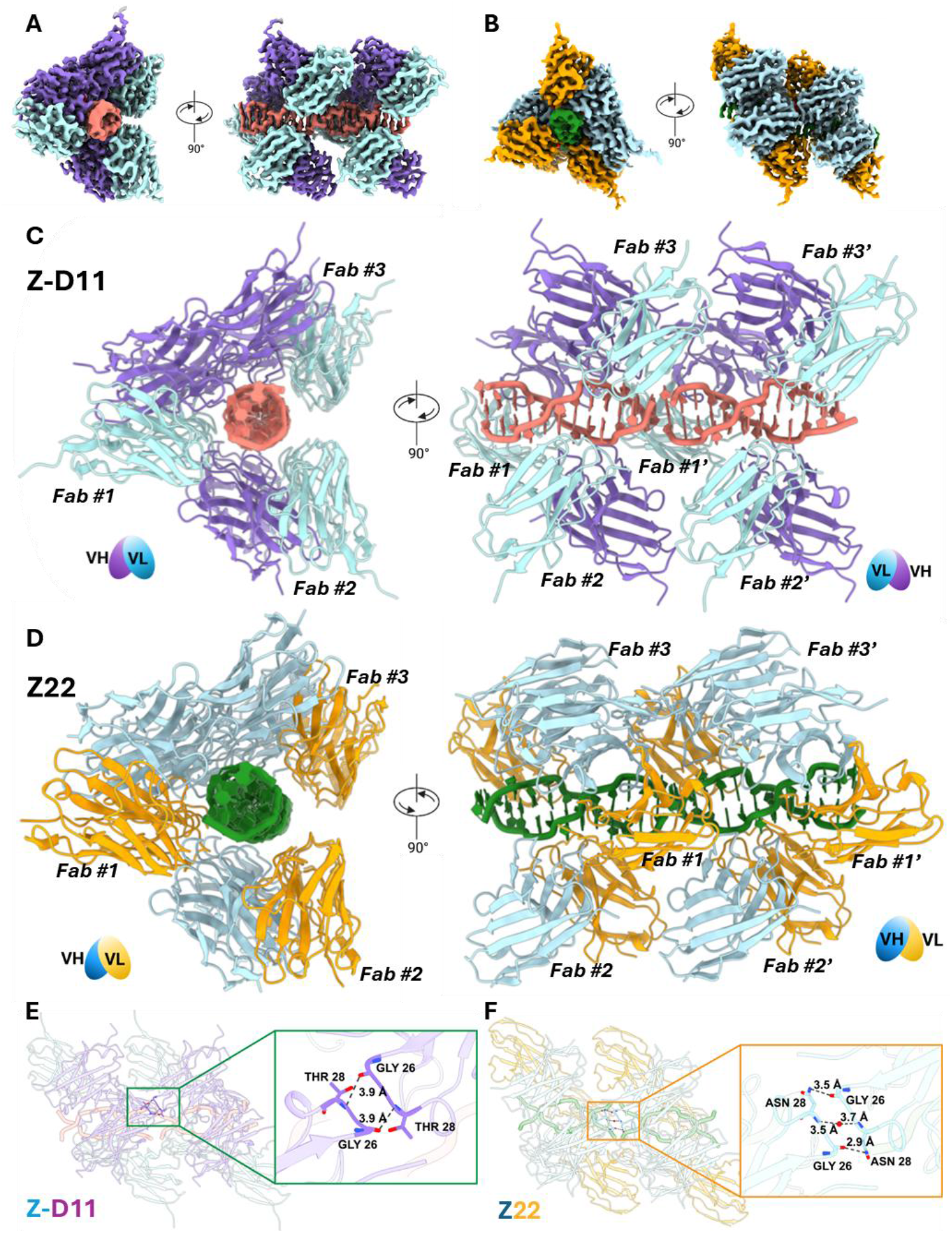
Cryo-EM structure of Z-DNA binding monoclonal antibodies to Z-DNA. Cryo-EM density map of **A**. Z-D11 mAb and **B**. Z22 mAb binding to two units of d(CG)_6_. Cryo-EM model of **C**. Z-D11 mAb-d(CG)_6_ and **D**. Z22 mAb-d(CG)_6_ complexes. Each fab within the trimer is labelled from 1 to 3, and the second trimer unit labelled from 1’ to 3’. Fab-fab interface of **E**. Z-D11 mAb-d(CG)_6_ and **F**. Z22 mAb-d(CG)_6_ complexes. Contacts between fab heavy chains are labelled with the corresponding bond lengths. Z-DNA is coloured in salmon and green for Z-D11 and Z22 complexes, respectively. Z-D11 is coloured in purple and turquoise for the heavy chain and light chain, respectively. Z22 is coloured in light blue and orange for the heavy chain and light chain, respectively.

### Molecular Basis for Z-DNA Recognition

The specificity of both Z-D11 and Z22 (mAbs) for the Z-form geometry is achieved through extensive contacts with both Z-form backbone strands and the guanine/cytosine bases accessible in the major groove. This recognition mechanism senses both the alternating *syn/anti* glycosidic bond configurations and the sugar pucker-mediated phosphate zig-zag geometry unique to Z-DNA. The Z-D11 Fab subunit aligns axially along the Z-DNA (Fig. 3A - 3D). Alignment of the individual fab subunits of one Z-D11 trimer shows no local conformational differences between the subunits, with root mean square deviations (RMSD) less than 0.35 across 123 atom pairs (Fig. 3A). The Z-D11 Fab engages the Z-DNA via Complementarity Determining Regions (CDRs) L1, L2, L3, H2, and H3 (Fig. 3B, 3D). For Z-D11, stabilization of the phosphate backbone involves charged residues, notably Arg 50 and Arg 52 (CDR-H2) and Arg 53 (CDR-L2), which form contacts ranging from 2.9 Å to 3.8 Å (Fig. 3C). The recognition mechanism of Z-D11 is further characterized by a strong presence of tyrosine residues, with Tyr 32 (CDR-L1) and Tyr 50 (CDR-L2) interacting directly with the exposed bases and sugar moieties in the major groove or via water-mediated contacts as seen with Tyr 105 and Tyr 106 on CDR-H3 (Fig 3D); Specific hydrogen bond contacts are formed by Tyr 105 and Tyr 106 at distances of 3.3 Å and 3.4 Å (Fig 3C).

**Figure 3.**
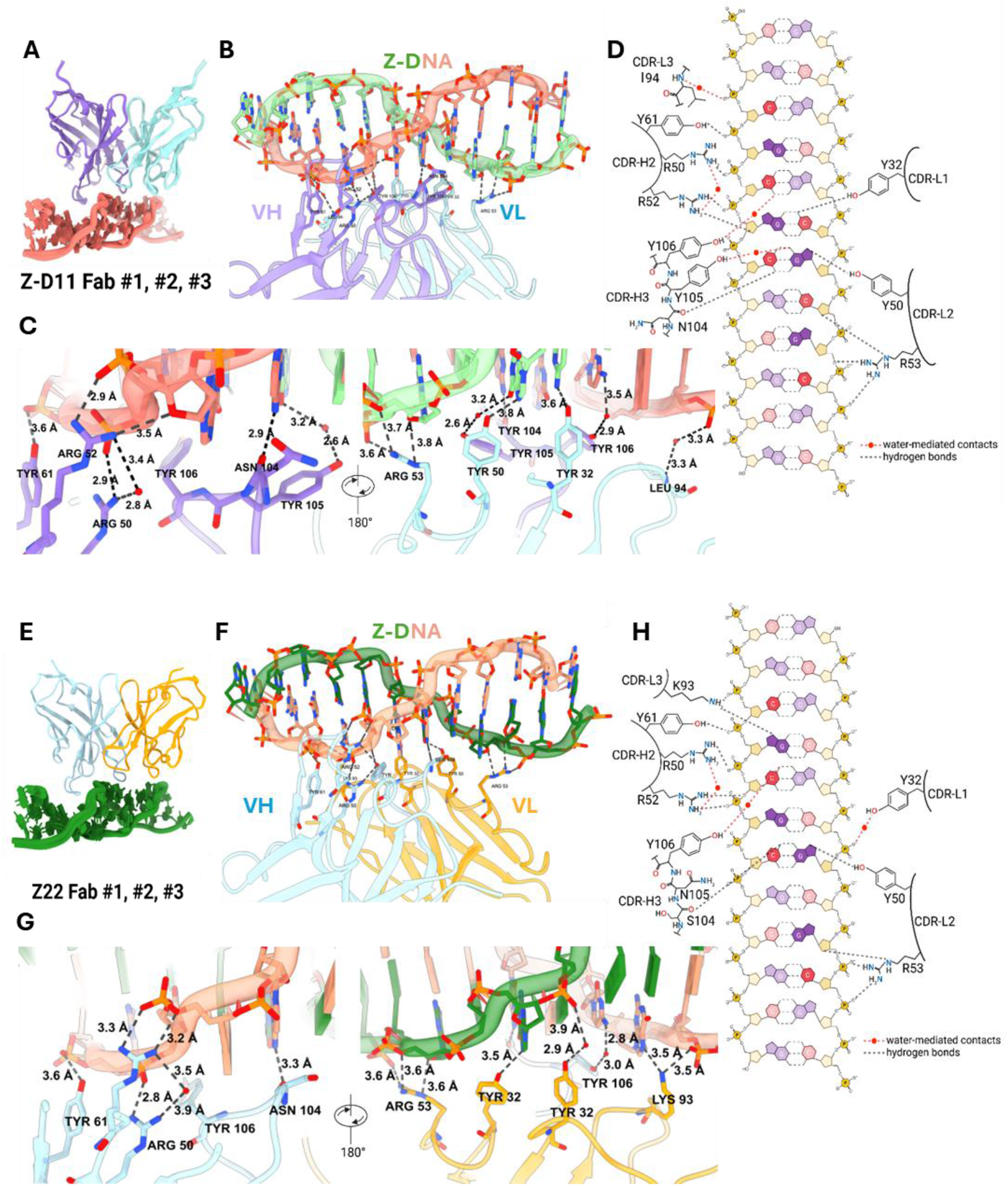
Z-D11 mAb and Z22 mAb interacts with Z-DNA via contacts along the sugar phosphate backbone and with the extruded bases. **A.** Alignment of the Z-D11 fab subunits along the d(CG)_6_ Z-DNA. **B. - C**. Interface of Z-D11 mAb binding to d(CG)_6_. **D**. Cartoon representation of the interaction between Z-D11 and d(CG)_6_. **E**. Alignment of the Z22 fab subunits along the d(CG)_6_ Z-DNA. **F. - G**. Interface of Z22 mAb binding to d(CG)_6_. **H**. Cartoon representation of the interaction between Z22 and d(CG)_6_. Each contact is labelled with its corresponding bond lengths. Each strand of Z-DNA is coloured in green and salmon separately, Z-D11 is coloured in purple and turquoise for the heavy chain and light chain, respectively. Z22 is coloured in light blue and orange for the heavy chain and light chain, respectively.

The Z22 Fab employs a similar axial recognition geometry (Fig 3E, 3H). Alignment of the individual fab subunits also shows no local conformational differences between the subunits, with root mean square deviations (RMSD) less than 0.25 across 121 atom pairs (Fig. 3E). Backbone contacts involve key residues such as Tyr 32 (CDR-L1), Arg 50, Arg 53 and Tyr 61 (CDR-H2), exhibiting bond lengths between 2.9 Å and 3.9 Å (Fig 3G). Base and sugar interactions in Z22 are facilitated by Tyr50, Arg 53 and Lys 93 from the light chain, and Ser 104 and Tyr 106 from the heavy chain (Fig. 3H).

### Conformational Selectivity Demonstrated by Molecular Modeling

To evaluate whether the recognition mechanisms employed by Z-D11 and Z22 are conformation-dependent rather than sequence-specific, we employed modeling of binding to both modified and alternative Z-DNA sequences (Fig. 4). First, modeling interactions with methylated Z-DNA (d^5^mCdG) demonstrated that both antibodies should maintain strong contacts; Z-D11 utilized key tyrosine residues (Tyr 106, Tyr 32, Tyr 105) with distances of 2.4–2.6 Å (Fig. 4A), while Z22 similarly maintained contacts through Tyr 106, Tyr 32, and Asn 105 (Fig. 4B). In fact, for Z-D11, detailed biochemical binding studies have revealed association constants of 60 pM with unmodified ds d(CG) oligo nucleotides in high salt conditions, while the d^5^mC modification weakened the affinity to 50 nM ^[35]^. Hence, while relative tight binding to (d^5^mCdG) as Z-conformer is retained with modified bases, the super tight binding to unmodified Z-DNA seems to be affected.

**Figure 4.**
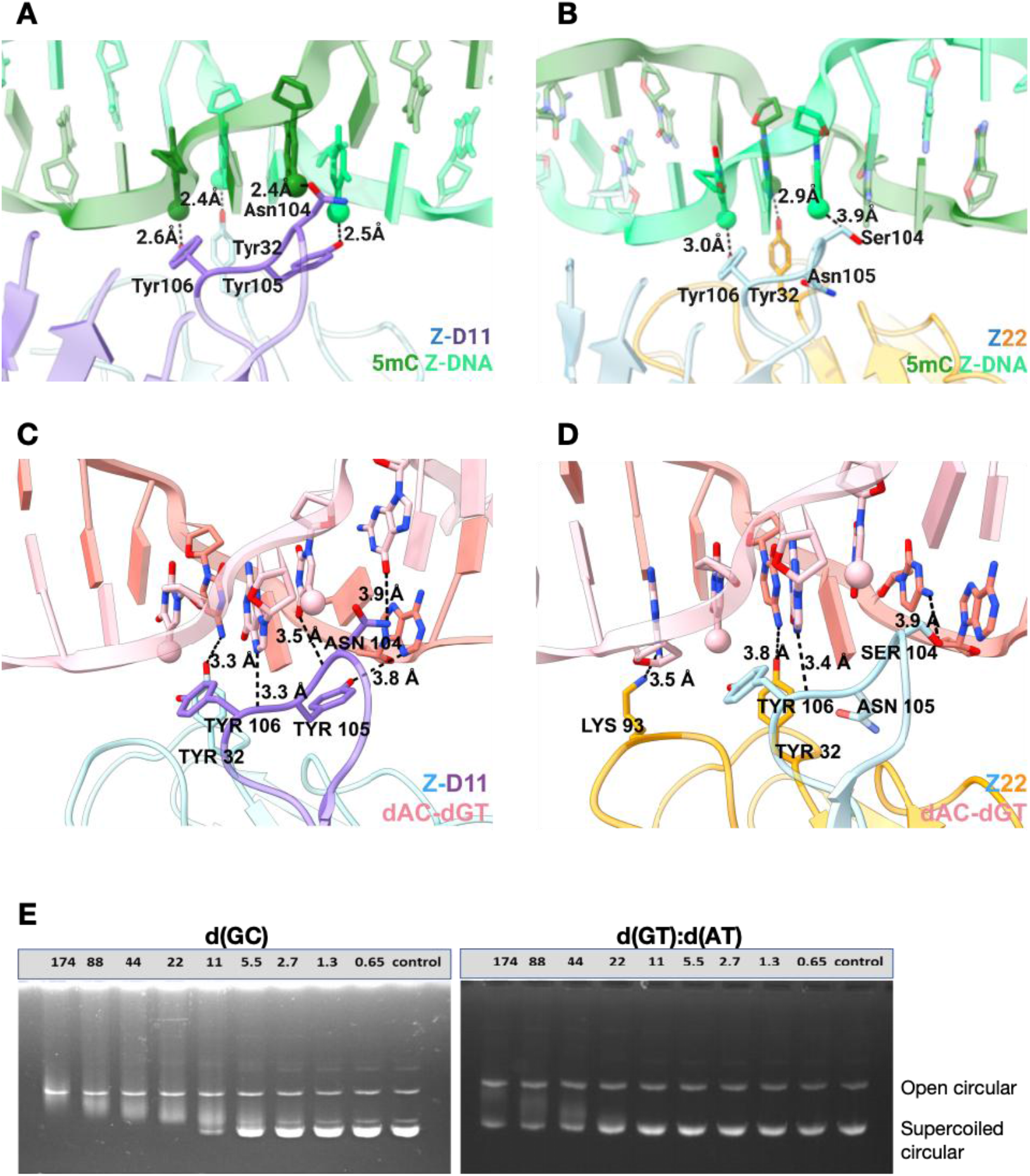
Modelling of Z-D11 and Z22 binding to alternative nucleic acids. Modelled interaction of **A**. Z-D11 mAb and **B**. Z22 mAb to 5mC Z-DNA. Modelled interaction interface of **C**. Z-D11 mAb and **D**. Z22 mAb to dAdC-dGdT Z-DNA. A cartoon representation is included above the binding interface, where the interacting nucleotides are highlighted. **E**. Electrophoretic Mobility Shift Assay (EMSA) of Z-D11 with plasmids containing sequences of d(GC)_12_ and d(GT):d(AT)_47_. Numbers represent the stoichiometric ratio of Z-D11:plasmid DNA. ^5^mC Z-DNA is coloured in green with the 5’-methyl group represented with a sphere, and dG nucleotides represented as a slab. dAdC-dGdT Z-DNA is coloured in salmon. Nucleotides not participating in the binding are represented as slabs. Z-D11 is coloured in purple and turquoise for the heavy chain and light chain, respectively. Z22 is coloured in light blue and orange for the heavy chain and light chain, respectively.

Second, modeling of binding to the sequence variant (dGdT:dAdC) indicated that binding capacity should be retained even with an altered purine/pyrimidine sequence: Z-D11 maintained backbone and base contacts via Tyr 32, Arg 53 and residues along the CDR-H3 (Fig. 4C), and Z22 retained significant interactions using multiple residues along multiple CDRs (Fig. 4D). For Z-D11, binding to (dGdT:dAdC) in Z-form under physiological conditions was confirmed experimentally using supercoiled plasmids and agarose EMSAs. Titration and comparison with binding to plasmids bearing a (dGdC:dGdC)_12_stretch in Z-form, however, indicated reduced affinity (Fig. 4E). Importantly, plasmids that do not contain the alternating purine/pyrimidine stretches in Z-conformation due to lack of supercoiling (open circular form), were not bound and shifted by Z-D11 during incubation, demonstrating that Z-D11 does not induce B-to-Z transitions under physiological conditions. These results collectively indicate that the dominant factor in recognition for both Z-D11 and Z22 is complementarity derived from Z-DNA’s unique geometry, hence suggesting their utility as sequence-agnostic conformation probes.

### Unique Recognition Mode for Left-handed Helices

Structural comparison of Z-D11 with established nucleic acid recognition antibodies, the dsRNA-binding antibody J2 ^[40]^ and the RNA:DNA hybrid-binding antibody S9.6 ^[41]^, revealed a shared architectural strategy: all three antibodies position the Complementarity Determining Regions (CDRs) loops to cradle the helical backbone and engage recurring geometric features of the ds-NA (Fig. 5A, 5B). Superposition of the Z-D11–Z-DNA complex with published J2–dsRNA and S9.6–RNA:DNA models showed that all three antibodies adopt a similarly angled approach toward the helix, positioning CDR-H2 and CDR-H3 to form the dominant contacts with the phosphate backbone, while CDR-L1 contributes stabilizing interactions at the minor-groove–facing surface (Fig. 5A, 5B). Multiple sequence alignments highlight that these key CDRs are conserved in overall length and loop type across the three antibodies, even though the identity of their contacting residues diverges substantially (Fig. 5A, 5B, S3A–B). Despite this shared framework, closer structural analysis of the CDR-dsNA interface coupled with multiple sequence alignment showed distinct mechanistic differences in molecular recognition, particularly within H2, H3, and L1 of the CDRs (Fig. 5A, 5B, S3A, SB). The recognition mechanism utilized by Z-D11 and Z22 differs fundamentally from these other mAbs by being specifically adapted to the Z-form’s left-handed geometry. While S9.6 recognizes hybrid asymmetry via 2′-OH and deoxyribose groups, and J2 recognizes A-form helices via backbone phosphate spacing, Z-D11 - and by extension, Z22 - sense the alternating *syn/anti* backbone torsions and the phosphate zig-zag geometry.

**Figure 5.**
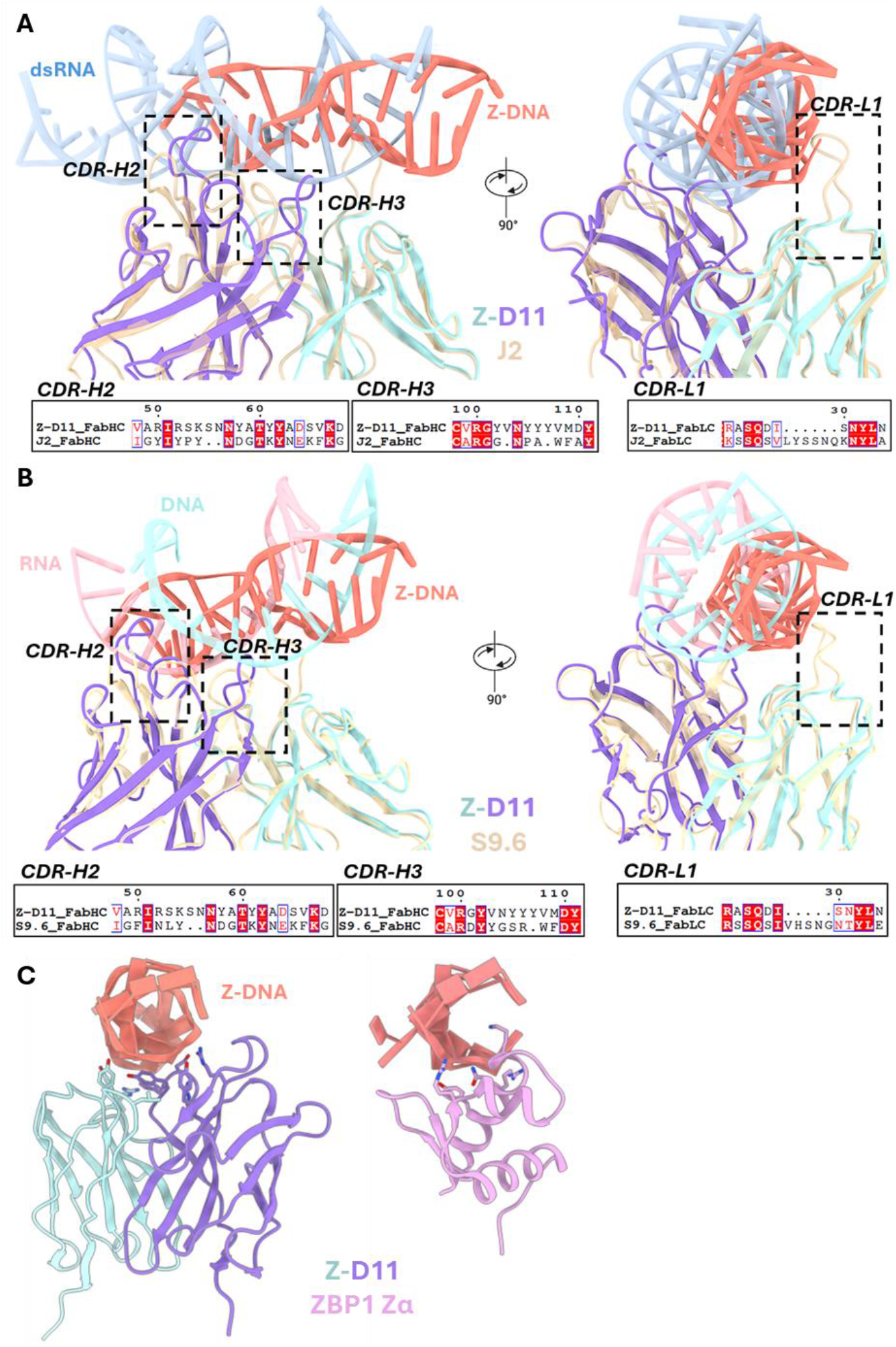
Comparison of nucleic acid binding mode of Z-D11 and Z22 with nucleic acid binding proteins. Overlay of Z-D11 with Z-DNA against **A**. dsRNA-binding antibody J2 with dsRNA and **B**. R-loop binding antibody S9.6 with DNA-RNA hybrid (PDB 7XLT). **C**. Side view comparison of Z-D11 against ZBP1 with Z-DNA (PDB 3EYI) down the Z-DNA axis. Z-DNA is coloured in salmon. dsRNA is coloured in medium blue; and DNA-RNA hybrid is coloured in light blue and pink. The differences between complementarity-determining regions (CDRs) are boxed with its corresponding amino acid sequences shown below. Z-D11 is coloured in purple and turquoise for the heavy chain and light chain, respectively. Both J2 and S9.6 are coloured in beige. ZBP1 Zα is coloured in pink.

To place these structural findings in a biological context, we further compared Z-D11 with the Zα domain of ZBP1^[42]^, a canonical Z-DNA sensor. Structural superposition shows that Z-D11 engages a face of the Z-helix distinct from the Zα-binding surface, approaching the DNA from a nearly orthogonal angle and contacting a non-overlapping stretch of backbone (Fig. 5C). Unlike Zα, which induces B-to-Z transitions and binds shallowly along one face of the helix, Z-D11 clamps the fully formed Z-helix from the opposite side through a wider, Fab-supported interface. This establishes that Z-D11 and Z22 detect pre-existing Z-DNA rather than promoting its formation, positioning them as orthogonal probes to endogenous Z-DNA–binding proteins.Together, these results demonstrate that Z-D11 and Z22 use a unique, geometry-driven recognition mode that distinguishes the Z-form from other nucleic acid conformations, while retaining the general CDR-based architectural logic seen in other dsNA-binding antibodies.

In order to further support our conclusions in a biological context, we performed immunofluorescent staining comparing Z-D11 and Z22 on CBL0137-induced Z-DNA formation in immortalized Zbp1−/− murine embryo fibroblasts (MEFs) ^[21, 43]^. The results revealed that Z-D11 stained nuclei of CBL0137-treated cells in a dose-dependent manner (Fig. 6A, 6B), manifesting a similar pattern as observed with Z22 (Fig. 6C, 6D). Nucleoli were not stained probably due to inaccessibility for mAb penetration rather than lack of Z-DNA formation. Furthermore, the complete absence of signals in CBL0137-untreated MEFs with either antibody demonstrates the specificity of these antibodies for Z-DNA in cellulo. They also suggest that detectable levels of Z-DNA, at least of longer Z-stretches, are not seen in these cells, and that neither mAb can induce B-to-Z transitions in fixed cells.

**Figure 6.**
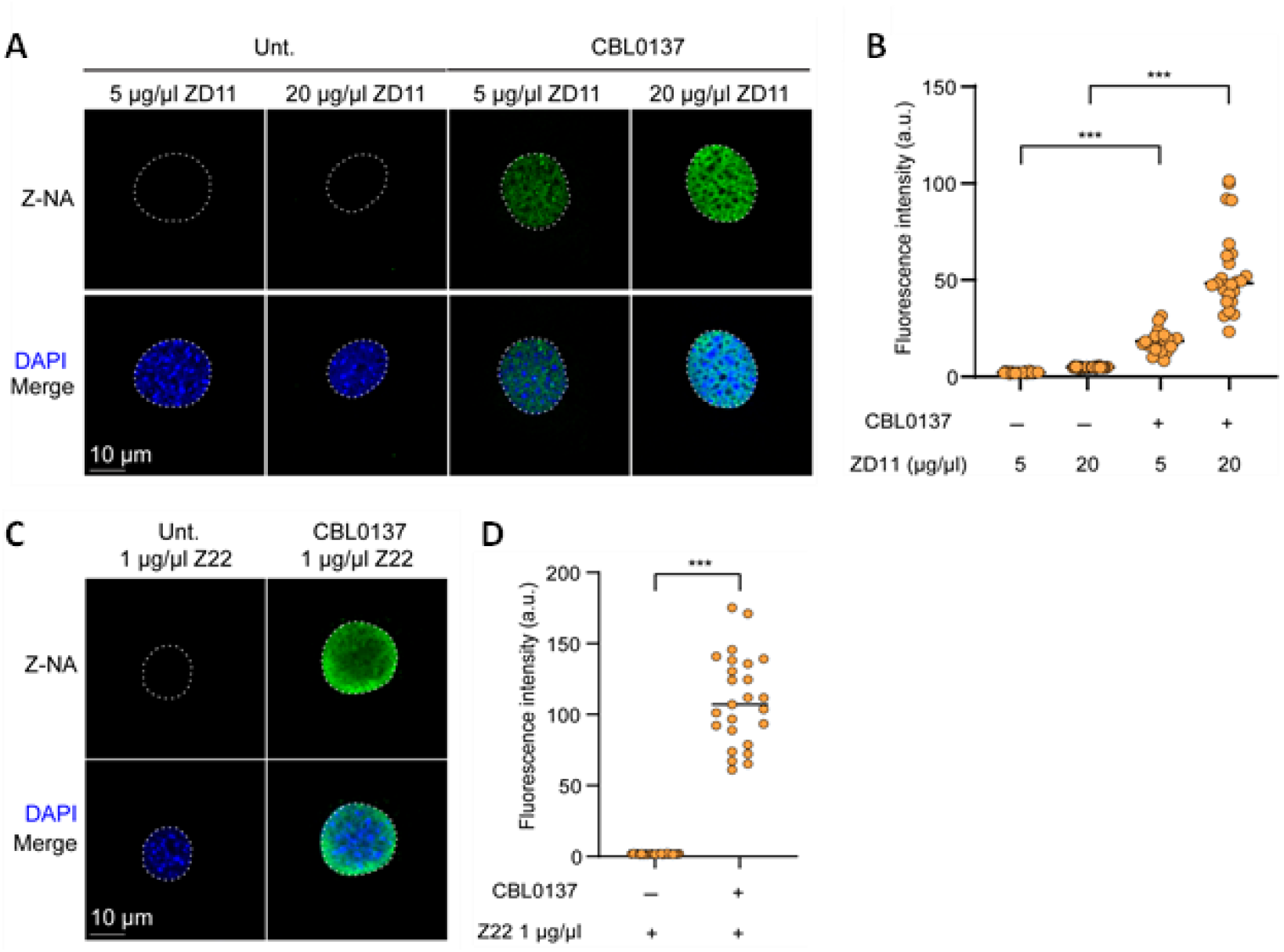
Comparison of Z-D11 and Z-22 Z-DNA staining in CBL0137-treated MEFs. **A**. Immortalized Zbp1-/- MEFs treated with CBL0137 (5µM) and fixed 6 h post-treatment were stained with mouse monoclonal anti-Z-NA antibody ZD11 at 5ug/ul or 20ug/ul, as indicated, and detected with Alexa488-conjugated secondary antibody (green). Nuclei are stained with DAPI (blue) and outlined with dashed lines. **B**. Quantification of fluorescence intensity of Z-NA signal shown in (**A**). **C**. Immortalized Zbp1-/- MEFs treated with CBL0137 (5µM) and fixed 6 h post-treatment were stained with rabbit monoclonal anti-Z-NA antibody Z22 at 1ug/ul, and detected with Alexa488-conjugated secondary antibody (green). Nuclei are stained with DAPI (blue) and outlined with dashed lines. **D**. Quantification of fluorescence intensity of Z-NA signal shown in (**C**). Data are mean ± SD. (n=25 cells/group). Data are representative of at least two independent experiments. Statistical significance was assessed using one-way analysis of variance (ANOVA) with Dunnett’s multiple-comparison test (**B**) and two-tailed unpaired t-tests with Welch’s correction (**D**) *** p < 0.0005. Scale bar reprensents 10 μm.

## Discussion

In this study, we reveal high-resolution cryo-EM structures of the Z-DNA–binding antibodies Z-D11 and Z22 in complex with ds d(CG)_6_ under Z-form favouring conditions, establishing their recognition principles at near-atomic detail. Both antibodies form highly ordered oligomeric assemblies around the left-handed duplex, driven by extensive interactions with the phosphate backbone, exposed bases in the major groove, and direct Fab–Fab contacts between the heavy chains. The filamentous arrangement observed for Z-D11, in particular, suggests that antibody–antibody interactions amplify avidity along extended Z-DNA tracts, contributing to super-tight binding and efficient detection even when Z-form segments are sparse or transient *in vivo*. These assemblies also highlight an unexpected architectural feature: Z-form recognition can be coupled to higher-order multimerization, a property that may enhance sensitivity while maintaining specificity for left-handed Z-DNA geometry.

Molecular modelling across alternative Z-form sequences reinforces that recognition by both antibodies is primarily governed by structural complementarity rather than base identity. The antibodies tolerate methylation and purine/pyrimidine substitutions while maintaining core interactions with the alternating syn/anti glycosidic torsions and the zig-zag phosphate topology characteristic of Z-form DNA. However, the Z-form sequence does affect binding affinity to some extent and should not be completely ignored [35]. Nevertheless, this substantial sequence-agnostic mode of recognition resolves long-standing ambiguity surrounding how Z-DNA antibodies distinguish the Z-form without enforcing a B-to-Z transition, and positions Z-D11 and Z22 as reliable probes that faithfully report endogenous Z-DNA rather than inducing it.

Comparison with canonical nucleic-acid-binding antibodies such as S9.6 (RNA:DNA hybrids) and J2 (A-form dsRNA) further clarifies the shared logic underlying structural epitope recognition: all three use CDR loops as geometric cradles that read recurring features of helical architecture. However, Z-D11 and Z22 diverge sharply in the type of geometry they detect. Instead of hybrid asymmetry or A-form spacing, these antibodies are specialized for the left-handed signatures of Z-DNA, establishing them as the first structurally resolved examples of antibodies adapted to a non-canonical nucleic acid conformation.

These findings have direct implications for understanding Z-DNA–binding proteins and the biological processes in which Z-form structures participate. Endogenous effectors such as ADAR1, ZBP1, and PKZ rely on Zα domains to sense the same geometric features captured by Z-D11 and Z22, and their activation depends on the transient formation of left-handed helices in chromatin or RNA-rich environments ^[6, 25, 44, 45]^. Z-D11 and Z22 offer a clearer and less perturbative readout of Z-DNA formation and localization since they do not induce B-to-Z transitions. The precise, sequence-independent detection of Z-form segments provides a much-needed experimental route to map where Z-DNA forms under physiological torsional stress, during transcription cycles, at CpG-rich promoters, and under inflammatory or antiviral signalling conditions. Their defined recognition footprint and high avidity for pre-existing Z-DNA make them well-suited for probing how structural transitions between B- and Z-forms contribute to gene regulation, chromatin remodelling, and immune responses. More broadly, these antibodies establish a framework for engineering future conformation-specific probes that can distinguish DNA and RNA states based solely on geometry. As such, Z-D11 and Z22 open new opportunities to interrogate the formation, regulation, and functional consequences of Z-DNA in living systems and provide a foundation for deeper mechanistic insight into the roles of left-handed nucleic acids across biology.

## Materials & Methods

### Cell Culture and Maintenance

Immortalized *Zbp1*^*−/−*^ murine embryo fibroblasts (MEFs) ^[43]^ were maintained in DMEM supplemented with 15% FBS, 1 mM sodium pyruvate, 1× GlutaMax and 1% penicillin– streptomycin, in a humidified incubator at 37°C, 5% CO_2_. Expi293 cells were maintained in OPM-293 media (OPM Biosciences) in a humidified shaking incubator at 37°C, 8% CO_2_.

### Protein Expression

PcDNA3.1 ZD11 IgG heavy chain and light chain plasmids were transfected in equal ratio into Expi293 cells directly using homemade polyethylenimine (PEI) at 1:3 DNA:PEI ratio. 18 hours post transfection, 5% (v/v) of OPM-293 Profeed (OPM Biosciences) was added to boost expression of the target proteins. Culture media was harvested five days post transfection.

### Protein Purification

In brief, harvested culture media was filtered and incubated with pre-equilibrated Protein G resin (GenScript) overnight on a roller platform at 4°C. The resin was collected in a gravity flow column and supernatant is passed through once more before washing the resin in 10 column volumes (CV) of wash buffer (1x PBS pH 7.4) twice. Antibodies were eluted from the column with 0.85 CV of elution buffer (0.1M glycine, pH 3), directly into a tube containing 15% of total volume of neutralization buffer (1M Tris, pH 8). Purified antibody was buffer exchanged into wash buffer and concentrated for downstream structural analysis.

### Cryo-EM sample preparation

Z22 mAb antibody was purchased from Absolute Antibody (cat # Ab00783-3.0). The 12-mer DNA, d(CG)_6_, was purchased from Integrated DNA Technologies (IDT). The sequence of d(CG)_6_is listed in Table 1. The DNA was resuspended in water to a final concentration of 2mM and annealed in a thermocycler using the following protocol: 95°C for 2 minutes, slow cooled to 4°C at a rate of 1°C/minute. For complexing under high salt conditions, annealed d(CG)_6_ was then diluted to 50µM in high salt buffer (20mM HEPES pH 7.4, 2.5M NaCl, 0.7M MgCl_2_) and incubated at 37°C for 30 minutes. The antibody was added in 3-fold molar excess to the DNA in high salt buffer. The resulting mixture was passed though analytical size exclusion chromatography using Superose 6 3.2/300 Increase column, pre-equilibrated in low salt buffer (20mM HEPES pH 7.4, 150mM NaCl). For complexing under low salt conditions, the antibody was mixed with the annealed d(CG)_6_ in low salt buffer. The mixture was subsequently passed though analytical size exclusion chromatography. Peak fractions containing the antibody and the DNA were collected for cryo-grid preparation.

| OGN | Length | Nucleic Acid Type | Sequence (5'-3') |
| --- | --- | --- | --- |
| d(CG)<sub>6</sub> | 12-mer | DNA | CGCGCGCGCGCG |

### Cryo-grid preparation

Cryo-grid preparation was performed with the Vitrobot Mark IV (Thermo Fisher Scientific). Cryo-grids used were 300-mesh gold grids with R1.2/1.3 holey carbon film (Quantifoil, EMS). The grids were glow discharged for 1 minute in air plasma immediately before use. 3µL sample was pipetted onto the grid surface, blotted for 8 to 12 seconds under blot force of 2 and immediately plunge frozen in liquid ethane. The grids were subsequently clipped for data collection.

### Cryo-EM data collection and processing

The dataset of Z-D11-d(CG)_6_ and Z22-d(CG)_6_ complexes were collected in-house (FACTS, NTU). Micrographs were recorded on the Titan Krios electron microscope operated at 300kV.

Detailed parameters for the data collection are summarized in Supplementary Table 1. The micrographs were motion corrected on CryoSPARC Live. The motion corrected micrographs were then imported into CryoSPARC-v4.5 for further processing.

For Z-D11-d(CG)_6_ sample, particles were extracted from 4,000 micrographs. An initial model was prepared and used for Topaz training and re-extraction before 2D classification. A total of 397,243 particles were manually selected from the 2D classes and separated into three classes by ab-initial reconstruction. These three classes were the subjected to heterogenous refinement, with one class showing clear densities that was selected for further processing. 147,917 particles from one 3D class were further refined to obtain a 2.67Å map. A final 2.55Å map with 146,988 particles of the Z-D11-d(CG)_6_ complex was obtained after reference-based motion correction and a final iteration of non-uniform and local refinements.

For Z22-d(CG)_6_ sample, particles were extracted from 4,527 micrographs. An initial model was prepared and used for Topaz training and re-extraction before 2D classification. 527,916 particles were manually selected from the 2D classes for multiple iterations of ab-initial reconstruction and heterogenous refinement. 60,600 particles from one class of heterogenous refinement was selected for reference-based motion correction and subjected to non-uniform refinement and local refinement to yield a 2.80Å map showing densities of two units of a Z22 trimer surrounding the Z-DNA. A mask was applied over the first trimer. Local refinement yielded a 2.67Å map with 59,879 particles showing the Z22 mAb in a trimer formation surrounding the Z-DNA.

The model of the antibody was prepared using AlphaFold 3. Each model was docked into the density map and manually refined using Coot and ChimeraX. The model was further refined in Phenix using real-space refinement with secondary structure and geometry restraints. All figures were generated by UCSF Chimera.

### EMSA

Binding assays were carried out as follows: about 250 ng of negatively supercoiled plasmids that carry either a (dG-dC)_12_ sequence stretch (top) or a (dG-dT)_47_ sequence stretch (bottom) in (partial) Z-conformation were incubated with increasing amounts of purified recombinant Z-D11 in standard PBS buffer for 60 minutes at room temperature. Samples without added mAb served as negative control. The reaction mixtures were subsequently loaded onto 1% agarose gels for electrophoresis in 0.5 x TBE buffer. Electrophoresis was for 2.5 hrs at 100 V, after which DNA was visualized by ethidium bromide staining under UV light using standard protocols. The (dG-dT) stretch was cloned from a mouse mast cell protease 1 promoter genomic sequence and B-to-Z-transitions in supercoiled plasmids were confirmed by two-dimensional gel electrophoresis ^[46]^.

### Immunofluorescence microscopy

CBL0137 (Catman Chemical Cat # 19110), Z-NA Ab Clone Z22 (Absolute Antibody, Cat # Ab00783-23.0), Alexa Fluor 488 donkey anti-rabbit IgG (H+L) (Invitrogen Cat # A21206), Alexa Fluor 488 donkey anti-mouse IgG (H+L) (Invitrogen Cat # A21202) were purchased from their indicated sources.

Cells were plated on 8-well glass slides (EMD Millipore) and allowed to adhere for at least 24 hours before use in experiments. Following treatment with CBL0137, cells were fixed for 10 min with freshly prepared 4% (w/v) paraformaldehyde in PBS, permeabilized in 0.5% (v/v) Triton X-100 in PBS, blocked with MAXblock blocking medium (Active Motif), and incubated overnight (16 h) with primary antibodies at 4 °C. After three washes in PBS, slides were incubated with fluorophore-conjugated secondary antibodies for 1 h at room temperature. Following an additional three washes in PBS, slides were mounted in ProLong Gold antifade reagent (Thermo Fisher Scientific) and imaged by Leica SP8 instrument. Fluorescence intensity was quantified using Leica LAS X software.

### Multiple Sequence Alignment

The sequences of Z22, J2 and S9.6 antibodies were obtained from published articles or deposited structures and aligned using ClustalOmega ^[47]^. Alignment was modified using ESPript ^[48]^ to obtain the structure-based alignment.

### Statistical Analysis

Statistical significance for immunofluorescence data was assessed using one-way analysis of variance (ANOVA) with either Dunnett’s multiple-comparison test or two-tailed unpaired t-tests with Welch’s correction. Data are presented as mean ± standard deviation. (n = 25 cells per group). Data is representative of at least two independent experiments. Statistical significance is indicated as p < 0.0005 (***).

## Acknowledgements

This research is supported by the Singapore Ministry of Education under its Singapore Ministry of Education Academic Research Fund Tier 3 (MOET32023-0003) and the Education Academic Research Funds Tier 1 (RT22/23 and RG84/21) to DL. Work in the SB lab was supported by NIH R01AI135025 and R01CA269975. We thank the scientific facility support from NTU Institute of Structural Biology and Protein Product Platform. We thank members of the DL lab for their support.

## Author Contributions

DHRC, PD and DL designed the study; DHRC, YL, YTL and PD performed the experiments; QV, ND participated in modeling strategies and discussions; ZH, CY, RMW, SB carried out immunofluorescence studies; all authors participated in data analysis; DHRC, YL, PD, and DL wrote the manuscript with inputs from all authors. DHRC and YL contributed equally to this work.

## Data And Code Availability

The SPA-cryoEM density map of the Z-D11-d(CG)_6_ complex has been deposited in EM Database under the accession code EMD-55905. The corresponding atomic coordinates have been deposited in the Protein Data Bank under accession code 9TGN. The SPA-cryoEM density map of the Z22-d(CG)_6_ dimer of trimer and trimer complex has been deposited in EM Database under the accession code EMD-55906 and EMD-55912, respectively. The corresponding atomic coordinates have been deposited in the Protein Data Bank under accession code 9TGO and 9TGW, respectively.

## References

1. Wang, A.H.J., et al., Molecular structure of a left-handed double helical DNA fragment at atomic resolution. Nature, 1979. 282(5740): p. 680–686.

2. Hall, K., et al., ‘Z-RNA’—a left-handed RNA double helix. Nature, 1984. 311(5986): p. 584–586.

3. Krall, J.B., et al., Structure and Formation of Z-DNA and Z-RNA. Molecules, 2023. 28(2).

4. Pohl, F.M. and T.M. Jovin, Salt-induced co-operative conformational change of a synthetic DNA: Equilibrium and kinetic studies with poly(dG-dC). Journal of Molecular Biology, 1972. 67(3): p. 375–396.

5. Rich, A. and S. Zhang, Timeline: Z-DNA: the long road to biological function. Nat Rev Genet, 2003. 4(7): p. 566–72.

6. Schwartz, T., et al., Crystal Structure of the Zα Domain of the Human Editing Enzyme ADAR1 Bound to Left-Handed Z-DNA. Science, 1999. 284(5421): p. 1841–1845.

7. Placido, D., et al., A left-handed RNA double helix bound by the Z alpha domain of the RNA-editing enzyme ADAR1. Structure, 2007. 15(4): p. 395–404.

8. Buzzo, J.R., et al., Z-form extracellular DNA is a structural component of the bacterial biofilm matrix. Cell, 2021. 184(23): p. 5740–5758.e17.

9. Fang, Y., et al., AIRE relies on Z-DNA to flag gene targets for thymic T cell tolerization. Nature, 2024. 628(8007): p. 400–407.

10. Yin, Q., et al., RPA1 protects DNA damage–induced PANoptosis in limb development. Science Advances, 2025. 11(34): p. eadw2756.

11. Qin, L.N., et al., Extrachromosomal DNA biogenesis is dependent on DNA looping and religation by YY1-Lig3-PARylation complex. Mol Cell, 2025. 85(16): p. 3090–3107 e11.

12. Song, Z., et al., Innate immune sensing of Z-nucleic acids by ZBP1-RIPK1 axis drives neuroinflammation in Alzheimer’s disease. Immunity, 2025. 58(10): p. 2574–2592.e9.

13. Pisetsky, D.S. and A. Herbert, The role of DNA in the pathogenesis of SLE: DNA as a molecular chameleon. Ann Rheum Dis, 2024. 83(7): p. 830–837.

14. Papatriantafyllou, M., Z-DNA as an inflammatory trigger in lupus. Nature Reviews Rheumatology, 2025. 21(6): p. 316–316.

15. Klein, B., et al., Epidermal ZBP1 stabilizes mitochondrial Z-DNA to drive UV-induced IFN signaling in autoimmune photosensitivity. Science Immunology, 2025. 10(105): p. eado1710.

16. Marshall, P.R., et al., Dynamic regulation of Z-DNA in the mouse prefrontal cortex by the RNA-editing enzyme Adar1 is required for fear extinction. Nat Neurosci, 2020. 23(6): p. 718–729.

17. Zhang, T., et al., Influenza Virus Z-RNAs Induce ZBP1-Mediated Necroptosis. Cell, 2020. 180(6): p. 1115–1129.e13.

18. Li, S., et al., SARS-CoV-2 Z-RNA activates the ZBP1-RIPK3 pathway to promote virus-induced inflammatory responses. Cell Res, 2023. 33(3): p. 201–214.

19. Yin, C., et al., Host cell Z-RNAs activate ZBP1 during virus infections. Nature, 2025.

20. DeAntoneo, C., A. Herbert, and S. Balachandran, Z-form nucleic acid-binding protein 1 (ZBP1) as a sensor of viral and cellular Z-RNAs: walking the razor’s edge. Current Opinion in Immunology, 2023. 83: p. 102347.

21. Zhang, T., et al., ADAR1 masks the cancer immunotherapeutic promise of ZBP1-driven necroptosis. Nature, 2022. 606(7914): p. 594–602.

22. Yang, Z.-H., et al., ZBP1 senses splicing aberration through Z-RNA to promote cell death. Molecular Cell, 2025. 85(9): p. 1775–1789.e5.

23. He, J., et al., ZBP1 senses spliceosome stress through Z-RNA:DNA hybrid recognition. Mol Cell, 2025. 85(9): p. 1790–1805.e7.

24. Jiang, X., et al., Spliceosome inhibition induces Z-RNA and ZBP1-driven cell death in small cell lung cancer. Cell Reports, 2025. 44(10).

25. Jiao, H., et al., Z-nucleic-acid sensing triggers ZBP1-dependent necroptosis and inflammation. Nature, 2020. 580(7803): p. 391–395.

26. Jiao, H., et al., ADAR1 averts fatal type I interferon induction by ZBP1. Nature, 2022. 607(7920): p. 776–783.

27. Herbert, A., et al., The Zalpha domain from human ADAR1 binds to the Z-DNA conformer of many different sequences. Nucleic Acids Res, 1998. 26(15): p. 3486–93.

28. Lushnikov, A.Y., et al., Interaction of the Zα domain of human ADAR1 with a negatively supercoiled plasmid visualized by atomic force microscopy. Nucleic Acids Research, 2004. 32(15): p. 4704–4712.

29. Lee, A.R., et al., Thermodynamic Model for B-Z Transition of DNA Induced by Z-DNA Binding Proteins. Molecules, 2018. 23(11).

30. Nichols, P.J., et al., Zα Domain of ADAR1 Binds to an A-Form-like Nucleic Acid Duplex with Low Micromolar Affinity. Biochemistry, 2024. 63(6): p. 777–787.

31. Brigido, M.M. and B.D. Stollar, Two induced anti-Z-DNA monoclonal antibodies use VH gene segments related to those of anti-DNA autoantibodies. J Immunol, 1991. 146(6): p. 2005–9.

32. Pohl, F.M., R. Thomae, and E. DiCapua, Antibodies to Z-DNA interact with form V DNA. Nature, 1982. 300(5892): p. 545–546.

33. Thomae, R., S. Beck, and F.M. Pohl, Isolation of Z-DNA-Containing Plasmids. Proceedings of the National Academy of Sciences of the United States of America, 1983. 80(18): p. 5550–5553.

34. Zarling, D.A., et al., Immunoglobulin recognition of synthetic and natural left-handed Z DNA conformations and sequences. J Mol Biol, 1984. 176(3): p. 369–415.

35. Thomae, R., Dissertation. 1984, University Konstanz, Konstanz, F.R.G.

36. Dröge, P., J.M. Sogo, and H. Stahl, Inhibition of DNA synthesis by aphidicolin induces supercoiling in simian virus 40 replicative intermediates. Embo j, 1985. 4(12): p. 3241–6.

37. Pohl, F.M., Hysteretic behaviour of a Z-DNA-antibody complex. Biophys Chem, 1987.26(2-3): p. 385–90.

38. Dröge, P. and A. Nordheim, Transcription-induced conformational change in a topologically closed DNA domain. Nucleic Acids Research, 1991. 19(11): p. 2941–2946.

39. Dröge, P. and F.M. Pohl, The influence of an alternate template conformation on elongating phage T7 RNA polymerase. Nucleic Acids Res, 1991. 19(19): p. 5301–6.

40. Bou-Nader, C., et al., Specificity and mechanism of the double-stranded RNA-specific J2 monoclonal antibody. bioRxiv, 2025.

41. Bou-Nader, C., et al., Structural basis of R-loop recognition by the S9.6 monoclonal antibody. Nat Commun, 2022. 13(1): p. 1641.

42. Ha, S.C., et al., The crystal structure of the second Z-DNA binding domain of human DAI (ZBP1) in complex with Z-DNA reveals an unusual binding mode to Z-DNA. Proceedings of the National Academy of Sciences, 2008. 105(52): p. 20671–20676.

43. Thapa, R.J., et al., DAI Senses Influenza A Virus Genomic RNA and Activates RIPK3-Dependent Cell Death. Cell Host Microbe, 2016. 20(5): p. 674–681.

44. Wu, C.X., et al., The Zalpha domain of PKZ from Carassius auratus can bind to d(GC)(n) in negative supercoils. Fish Shellfish Immunol, 2010. 28(5-6): p. 783–8.

45. Herbert, A., Z-DNA and Z-RNA in human disease. Communications Biology, 2019. 2(1):p. 7.

46. Kulish, V.V., L. Heng, and P. Dröge, Z-DNA-induced super-transport of energy within genomes. Physica A: Statistical Mechanics and its Applications, 2007. 384(2): p. 733–738.

47. Sievers, F., et al., Fast, scalable generation of high-quality protein multiple sequence alignments using Clustal Omega. Mol Syst Biol, 2011. 7: p. 539.

48. Robert, X. and P. Gouet, Deciphering key features in protein structures with the new ENDscript server. Nucleic Acids Research, 2014. 42(W1): p. W320–W324.

