## Supplementary Information for "Cryo-EM structures of anti Z-DNA antibodies in complex with antigen reveal distinct recognition modes of a left-handed geometry"

Z-DNA, left-handed geometry; monoclonal antibody, cryo-EM structure, antibody avidity

**Supplementary Materials**

Supplementary Figures S1-S3

Supplementary Table 1

**Supplementary Figure S1. Structure determination of the Z-D11-d(CG)$_6$-DNA complex. A.** Workflow for the determination of the structure Z-D11-d(CG)$_6$. **B.** Angular distribution of particles used for the final 3D reconstruction. **C.** Local resolution distribution of the final map. **D.** FSC curves of non-uniform 3D refinement obtained from cryoSPARC.

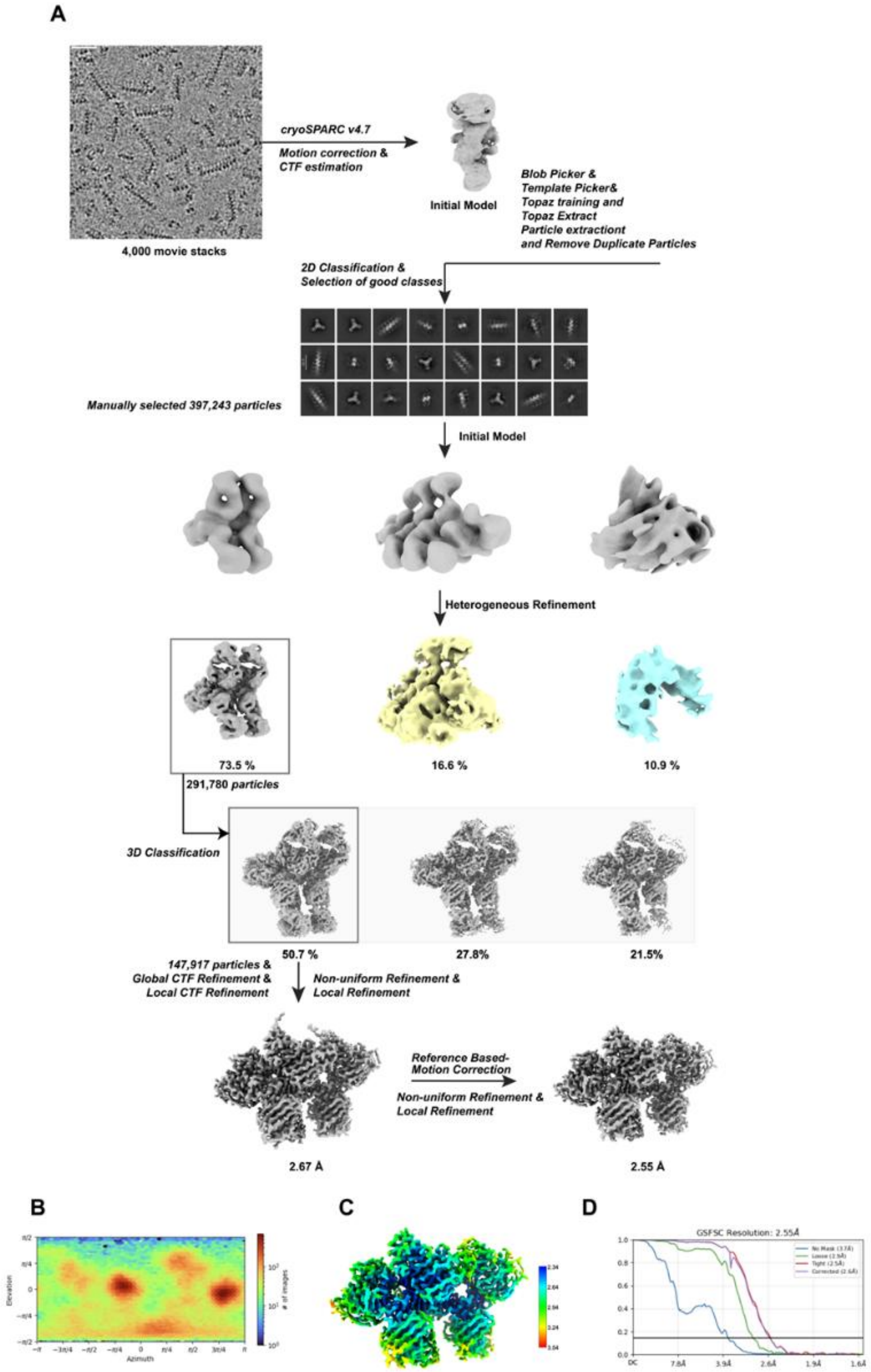

**Supplementary Figure S2. Structure determination of the Z22-d(CG)$_6$-DNA complex. A.** Workflow for the determination of the structure Z22-d(CG)$_6$. **B.** Angular distribution of particles used for the final 3D reconstruction. **C.** Local resolution distribution of the final map. **D.** FSC curves of non-uniform 3D refinement obtained from cryoSPARC.

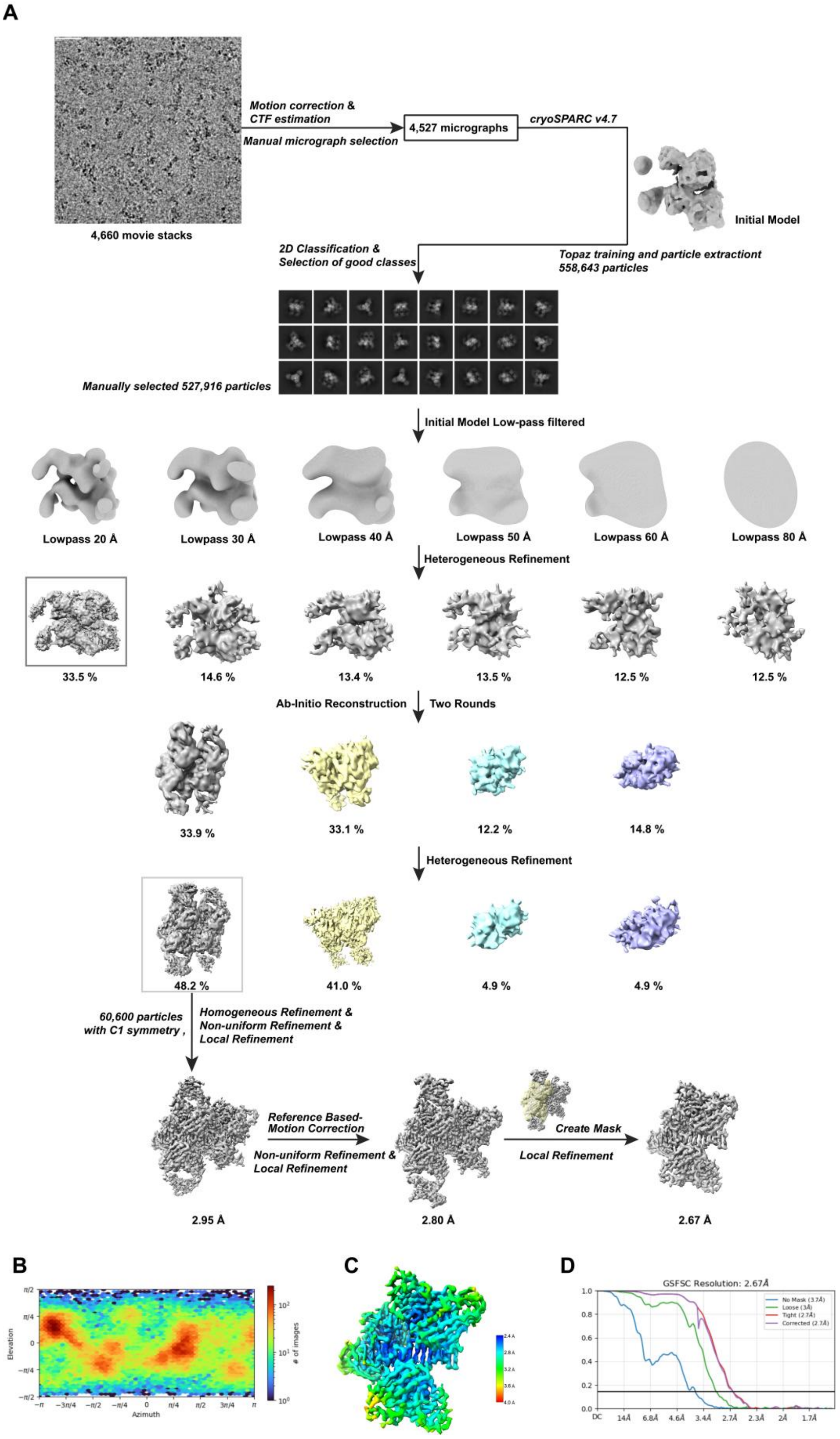

**Supplementary Figure S3. Multiple sequence alignment (MSA) of Z-DNA binding antibodies Z-D11 and Z22 fragment antigen-binding (fab) region with R-loop binding antibody S9.6 and dsRNA binding antibody J2.** MSA of the **A.** fab light chain, and **B.** fab heavy chain. The complementarity determining region (CDR) is underlined following the Kabat/Chothia numbering scheme. Alignment is based on Z-D11 antibody. MSA prepared using ClustalOmega.

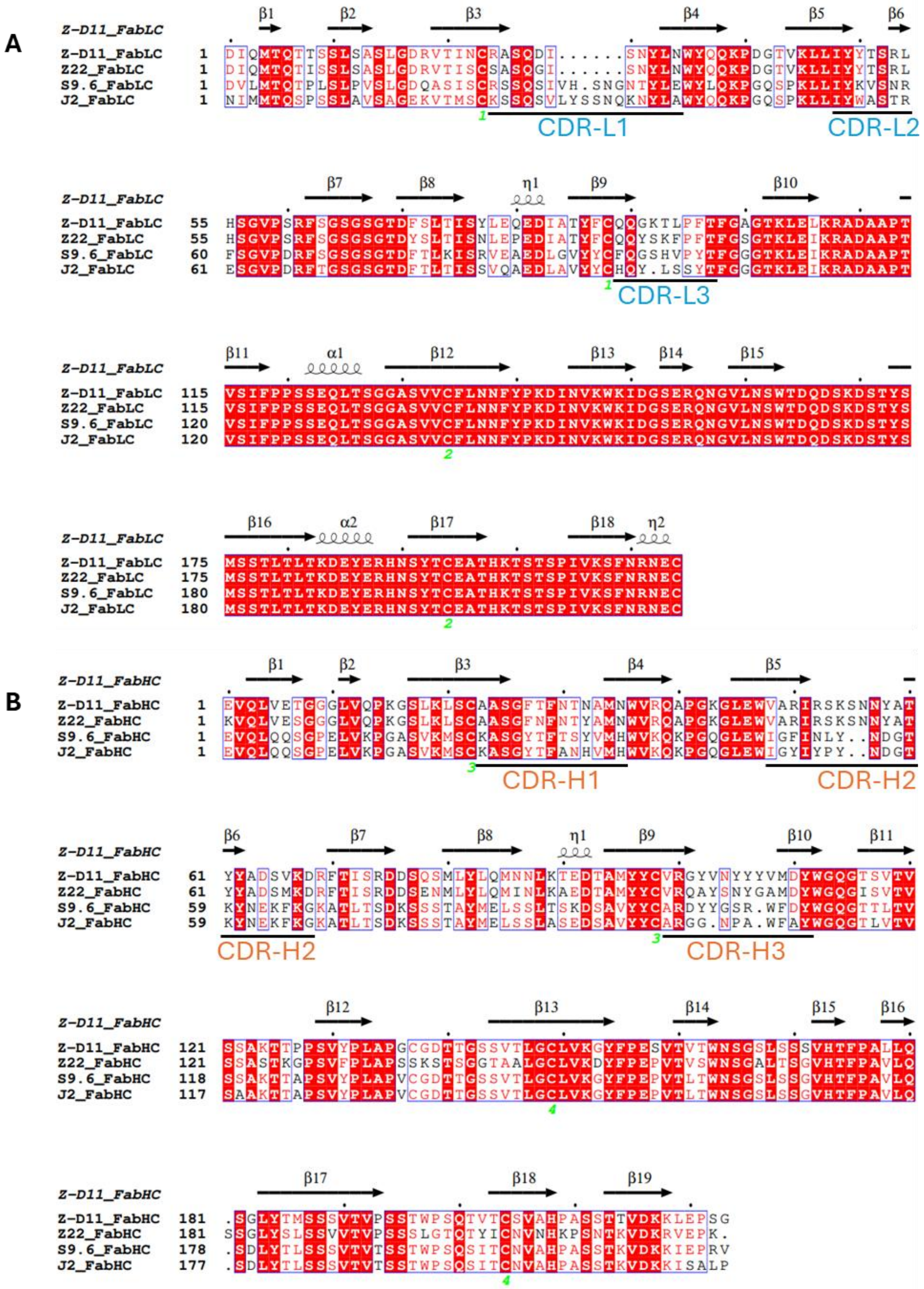

**Supplementary Table 1. Collection, refinement and validation statistics for cryo-EM density map and model.** Collection details for the cryo-EM map presented in the text are shown above, and validation statistics for the model generated from the map are shown below.

| | **Z-D11-d(CG)$_6$ (dimer of trimer)** | **Z22-d(CG)$_6$ (dimer of trimer)** | **Z22-d(CG)$_6$ (trimer)** |
|---|---|---|---|
| **EMDB:** | 55905 | 55906 | 55912 |
| **PDB:** | 9TGN | 9TGO | 9TGW |
| **Data collection and processing** | | | |
| Detector | Falcon 4i | | |
| Magnification | 165,000 × | | |
| Voltage (kV) | 300 | | |
| Electron exposure (e–/Å$^2$) | 50 | | |
| Defocus range (μm) | -0.5 ~ -1.5 | | |
| Pixel size (Å) | 0.76 | | |
| Symmetry imposed | C1 | C1 | C1 |
| Initial particle images (no.) | 494,925 | 558,643 | |
| Final particle images (no.) | 146,988 | 59,879 | 59,879 |
| Map resolution (Å) | 2.55 | 2.80 | 2.67 |
| FSC threshold micrographs | 0.143 | 0.143 | 0.143 |
| Map resolution range (Å) | 2.31 – 2.91 | 2.58 – 3.13 | 2.39 - 3.00 |
| **Refinement** | | | |
| Initial model used (PDB code) | AlphaFold | | |
| Model resolution (Å) | 3.0 | 1.8 | 2.0 |
| FSC threshold | 0.143 - 0.5 | | |
| Model resolution range (Å) | 2.5 - 3.0 | 1.5 - 2.9 | 1.5 ~ 3.1 |
| Map sharpening B-factor (Å$^2$) | 57.4 | 58.7 | 66.3 |
| **Model composition** | | | |
| Non-hydrogen atoms | 11940 | 11676 | 5830 |

| | | | |
|---|---|---|---|
| Protein residues | 1392 | 1362 | 681 |
| Nucleotide | 48 | 48 | 24 |
| Water | 22 | 32 | 15 |
| Ligands | 8 (Mg) | 15 (Mg) | 0 |
| **B factors (Å$^2$)** | | | |
| Protein | 5.06/130.01/60.48 | 10.51/130.67/59.34 | 17.21/90.77/52.02 |
| Nucleotide | 11.24/70.98/32.47 | 13.41/109.65/35.68 | 6.41/78.02/29.40 |
| Water | 23.01/61.95/37.12 | 9.29/85.84/44.84 | 28.88/62.03/46.62 |
| Ligand | 33.35/52.85/41.84 | 38.66/54.17/47.29 | N/A |
| **R.M.S. Deviations** | | | |
| Bonds length (Å) | 0.005 (1) | 0.003 (0) | 0.002 (0) |
| Bonds Angle (˚) | 0.690 (5) | 0.599 (0) | 0.499 (0) |
| **Validation** | | | |
| MolProbity score | 1.94 | 1.64 | 1.61 |
| Clashscore | 3.60 | 4.28 | 4.01 |
| Rotamer outliers (%) | 5.28 | 3.01 | 3.61 |
| **Ramachandran plot** | | | |
| Favored (%) | 96.35 | 97.68 | 98.06 |
| Allowed (%) | 3.65 | 2.17 | 1.94 |
| Disallowed (%) | 0.00 | 0.15 | 0.00 |
